# Open Targets Genetics: An open approach to systematically prioritize causal variants and genes at all published human GWAS trait-associated loci

**DOI:** 10.1101/2020.09.16.299271

**Authors:** Edward Mountjoy, Ellen M. Schmidt, Miguel Carmona, Gareth Peat, Alfredo Miranda, Luca Fumis, James Hayhurst, Annalisa Buniello, Jeremy Schwartzentruber, Mohd Anisul Karim, Daniel Wright, Andrew Hercules, Eliseo Papa, Eric Fauman, Jeffrey C. Barrett, John A. Todd, David Ochoa, Ian Dunham, Maya Ghoussaini

## Abstract

Genome-wide association studies (GWAS) have identified many variants robustly associated with complex traits but identifying the gene(s) mediating such associations is a major challenge. Here we present an open resource that provides systematic fine-mapping and protein-coding gene prioritization across 133,441 published human GWAS loci. We integrate diverse data sources, including genetics (from GWAS Catalog and UK Biobank) as well as transcriptomic, proteomic and epigenomic data across many tissues and cell types. We also provide systematic disease-disease and disease-molecular trait colocalization results across 92 cell types and tissues and identify 729 loci fine-mapped to a single coding causal variant and colocalized with a single gene. We trained a machine learning model using the fine mapped genetics and functional genomics data using 445 gold standard curated GWAS loci to distinguish causal genes from background genes at the same loci, outperforming a naive distance based model. Genes prioritized by our model are enriched for known approved drug targets (OR = 8.1, 95% CI: [5.7, 11.5]). These results will be regularly updated and are publicly available through a web portal, Open Targets Genetics (OTG, http://genetics.opentargets.org), enabling users to easily prioritize genes at disease-associated loci and assess their potential as drug targets.

## Introduction

Over 90% of GWAS-associated SNPs fall in non-coding regions, indicating that they affect expression of neighbouring genes through regulatory mechanisms ^1,2^, which can act over long distances and affect more than one gene. Hence, identification of the causal gene(s) and cell or tissue site of action is a major challenge requiring detailed low-throughput analysis of individual loci. One default approach has been to assign the top trait-associated SNP to the closest gene at each locus. However relying on physical proximity alone can be misleading since SNPs can influence gene expression over long genomic ranges ^3^, with studies based on eQTL data suggesting that two thirds of the causal genes at GWAS loci are not the closest ^4,5^. To add to the challenge, associated SNPs often span large regions due to linkage disequilibrium (LD), and pinning down the functional SNP and the tissue or cell type which mediates its effect can be complicated.

Connecting causal variants with their likely causal gene is a laborious process which requires the integration of GWAS data with multi-omics datasets across a wide range of cell types and tissues such as expression and protein quantitative traits (eQTL and pQTL), chromatin accessibility and chromatin interaction datasets. Subsequent functional assessment (such as reporter assays and CRISPR/Cas9 genome editing) can then be used to confirm the relationship between a putative causal variant and the gene it regulates. Using these integrative approaches, systematic international efforts have been undertaken to translate GWAS associated signals into target genes focused on one or a small subset of phenotypes ^6–9^. However, there are currently no resources that systematically prioritize all genes beyond specific therapy areas ^9^. Therefore, there is a need for a comprehensive, unbiased, scalable and reproducible approach that leverages all the publicly available data and knowledge to assign genes systematically to published loci across the entire range of phenotypes and diseases.

Drug development is hindered by a high attrition rate, with over 90% of the drugs that enter clinical trials failing, primarily due to lack of efficacy found in later, more costly stages of development ^10^. Retrospective analyses have estimated that drugs are twice as likely to be approved for clinical use if their target is supported by underlying GWAS evidence ^11^. Hence there is a critical need to build strategies that incorporate novel genetic discoveries and mechanistic evidence from GWAS and post-GWAS studies to suggest novel therapeutic targets for which to develop medicines, and ultimately increase the success rate of drug development.

Here, we describe a universal solution to these challenges: a systematic and comprehensive analysis pipeline for integrating GWAS results with functional genomics data to prioritize the causal gene(s) at each published GWAS-associated locus. The pipeline performs fine-mapping and systematic disease-disease and disease-molecular trait colocalization analysis. We integrate information from GWAS, expression and protein quantitative trait loci (eQTL and pQTL) and epigenomics data (e.g. promoter capture Hi-C, DNase hypersensitivity sites). For gene prioritization we developed a machine learning model trained on a set of 445 curated gold-standard GWAS loci for which we have moderate or strong confidence in the functionally implicated gene. The model integrates the fine-mapping with the functional genomics data, gene distance, and in silico functional predictions to link each locus to its target gene(s). This output of this pipeline feeds into Open Targets Genetics (https://genetics.opentargets.org), a user-friendly, freely available, integrative web portal enabling users to easily prioritize likely causal variants and target genes at all loci and assess their potential as pharmaceutical targets through linking out to Open Targets Platform ^12,13^ and will be regularly updated as new data become available.

## Results

### Pipeline Overview

We harmonised and processed GWAS data from the GWAS Catalog and from UK Biobank, and conducted systematic fine mapping to generate sets of credibly causal variants across all 133,441 study-lead variant associated loci. We also conducted cross-trait colocalization analyses for 3,621 GWAS studies with summary statistics available, which enabled us to identify traits and diseases that share common genetic etiology and mechanisms. To investigate whether changes in gene expression and protein abundance influence trait variation and disease susceptibility, we integrated 92 tissue- and cell type-specific molecular QTL datasets including GTEx ^14^, eQTLGen ^15^, the eQTL Catalogue ^16^ and pQTLs ^17^ and conducted systematic disease-molecular trait colocalization tests. Finally, we used a machine learning framework based on fine mapping, colocalization, functional genomics data and distance to prioritize likely causal genes at all trait-associated loci (Figure 1).

**Figure 1:**
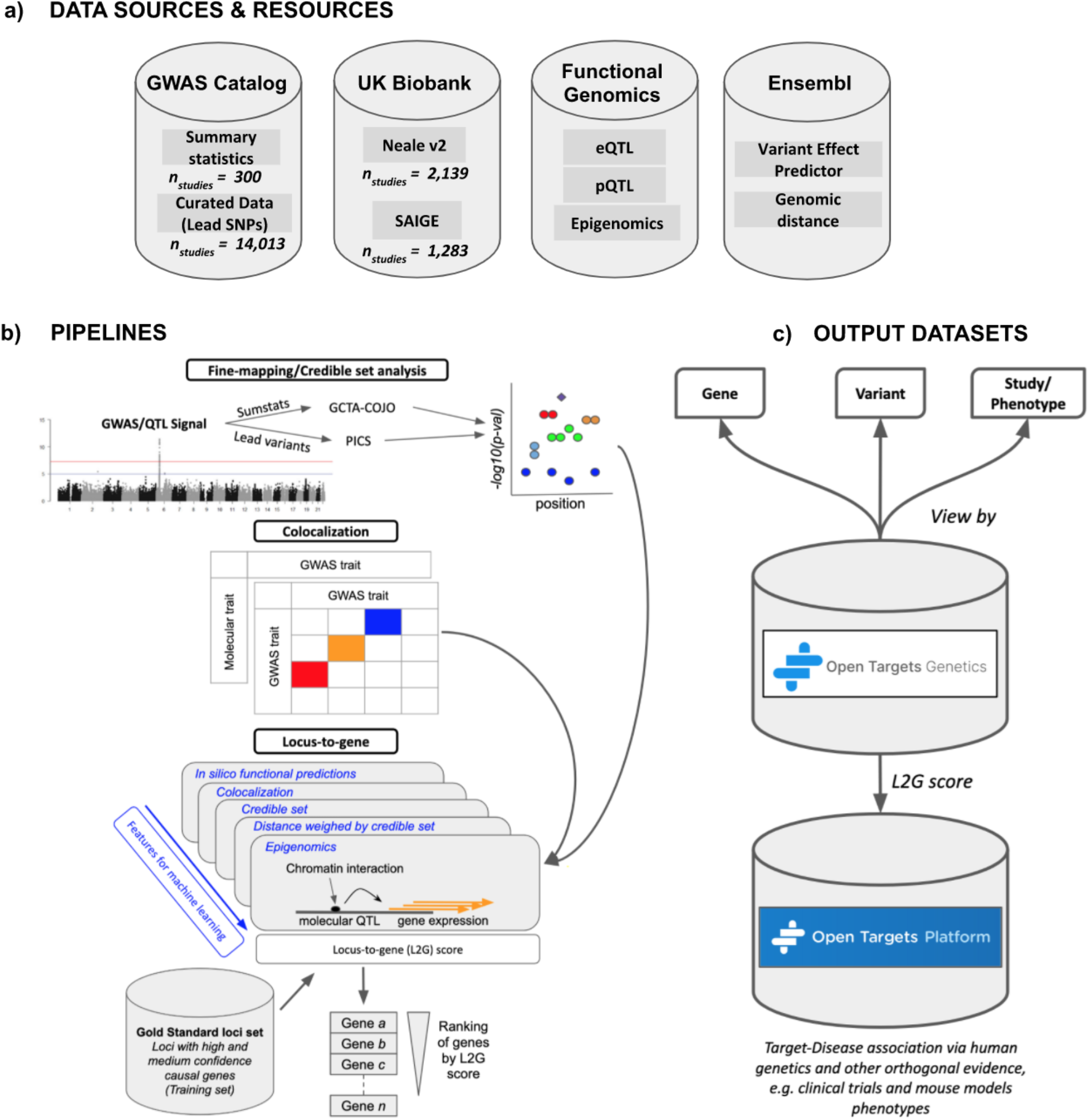
Open Targets Genetics pipeline schematic. a) Data sources include all available GWAS, as well as variant effect predictions and functional genomic data. b) A number of pipelines are run to perform statistical fine-mapping of GWAS, colocalization with gene expression quantitative trait studies (QTLs) and also between distinct GWAS traits, and integrative “locus-to-gene” prioritization from both genetic and functional genomic input features. c) Outputs of the pipelines are available in a web portal, via programmatic API, and as bulk downloads.

### Fine mapping of all published genome-wide association studies

To establish a comprehensive resource linking variants and traits or diseases, we integrate GWAS studies both with and without full summary statistics. Full summary statistics were obtained from three sources: the NHGRI-EBI GWAS Catalog summary statistics database (number of studies (n_study_) = 300)^18^; binary phenotypes from UK Biobank as published by Zhou et al. (n_study_= 1,283) ^19^ and all other UK Biobank phenotypes from the Neale lab (n_study_= 2,139; downloaded 21/01/2019)^20^ Studies with full summary statistics were restricted to those of predominantly European ancestries due to the lack of suitable reference genotypes required for conditional analysis from other populations. Studies without full summary statistics included all others in the NHGRI-EBI GWAS Catalog (n_study_= 14,013)^18^. To prioritize candidate causal variants at each GWAS association, we performed fine mapping of 10,494 GWAS Catalog and UK Biobank studies. Two fine-mapping methods were used to maximise coverage of GWAS studies, one using full summary statistics and a second using linkage disequilibrium (LD) information only (see methods). For studies with full summary statistics, we first identified independent signals using GCTA-COJO ^21^ and then conducted per-signal conditional analysis adjusting for other independent signals in a region ±2 Mb from the sentinel variant. We then used the Approximate Bayes Factor approach ^22^ to fine-map each conditionally independent signal. For studies without summary statistics, we used the PICS method ^23^ with an LD reference from the most closely matched 1000 Genomes superpopulation to estimate the probability that each variant is causal. Both methods output a posterior probability (PP) for each variant to be causal for the given association.

A total of 133,441 sentinel variants were detected, with 53% of these being shared by more than one study (70,860 distinct sentinel variants). To assess the concordance of the two methods we compared the 95% credible sets after applying both methods to all loci from studies with summary statistics available. We found a median absolute difference in credible set size of 7 variants (Supplementary Figure 1a), whereas the median credible set contained 17 variants. On average across loci, 70% of the credible set posterior probability colocated to the same variants between the two methods (Supplementary Figure 1b). These results suggest that on average the methods produced have comparable results. For subsequent analyses, we therefore used the full summary statistics method where these data were available, and for studies without summary statistics we used the PICS method.

Out of 133,441 loci association signals, 12,500 (9%) could be resolved to a single variant having PP > 0.95 and a further 21,279 (16%) to between 2 and 5 likely causal variants. Single-variant credible sets were 8.5 times more likely to have a moderate or high impact on protein-coding transcripts as predicted by the Ensembl variant effect predictor (VEP) ^24^ compared to variants in credible sets with 2 or more variants (OR=8.51, p<2.2e^−16^, Fisher’s exact test). Outside coding regions, single-variant credible set variants were preferentially located in Ensembl Regulatory Build regulatory elements, including: promoters (OR=1.70, p<2.2e^−16^), enhancers (OR=1.09, p=4.08e^−4^), transcription factor binding motifs (OR=1.85, p=1.22e^−15^) or other open chromatin regions (OR=1.19, p=4.8e^−5^).

In order to identify GWAS signals with high-confidence evidence linking the trait to variant and variant to gene, we took single-variant resolution loci and filtered these to retain variants with moderate or high-impact coding consequences in VEP. We identified 2,284 single coding variants linking 378 genes to 303 traits (Supplementary Table 1). Among these were several known disease-causal gene associations and targets of approved therapies (Supplementary Table 2) as well as novel disease-causal gene associations that had no prior evidence in the Open Targets Platform. One example is rs35383942, associated with breast cancer ^19,25^, which is a predicted deleterious missense variant (Arg28Gln, CADD=24.3) in *PHLDA3* (Pleckstrin Homology Like Domain Family A Member 3). PHLDA3 is the direct target of TP53 and acts as a tumor suppressor gene through inhibition of AKT1, an oncogene that plays a pivotal role in cell proliferation and survival ^26^.

### Colocalization of GWAS and molecular traits

Since most associated variants are non-coding, it is expected that they influence disease risk through alteration in gene expression or splicing. One way to identify the target gene is to demonstrate that the statistical association of a GWAS locus and a gene expression QTL are colocalized -- that is, that the pattern of SNP associations is consistent with them sharing the same causal variant. We conducted systematic colocalization analysis ^27^ of GWAS loci with molecular trait QTLs from 92 tissues or cell types. The QTL datasets (Supplementary Table 3) include pQTLs for 2,994 plasma proteins assessed in 3,301 individuals of European descent ^17^, eQTLs from 48 GTEx tissues (v7.0), blood eQTLGen ^15^, and 14 eQTL studies from the newly established eQTL Catalogue, a resource of uniformly processed gene expression and splicing QTLs recomputed from previously published datasets ^16^. The results of the colocalization test are summarised by the probability, referred to as “H4”, that a causal variant is shared.

GWAS-molecular QTL loci were tested if there was at least 1 variant overlapping in their 95% credible sets, suggesting prior evidence for colocalization (refer to methods). Of the 70,364 trait-associated loci from studies with summary statistics available, 49.4% had no colocalizing gene at an H4 threshold >0.8, 25.5% had exactly 1 colocalizing gene and 25.2% had >1 colocalizing gene. For loci with evidence of colocalization between GWAS and molecular QTL traits, 29% were specific to a single tissue or cell type, whereas 71% were observed across multiple tissues. We also examined non-coding QTLs that were fine-mapped to a single-variant resolution, and which colocalized with binary traits GWAS (H4>0.95). Results from this analysis are summarised in Supplementary Table 4.

We also performed cross-trait colocalization across 3,621 GWAS to identify traits that are likely to be underpinned by the same molecular mechanism. A summary of the binary trait GWAS loci with the highest colocalization score (H4>0.95) is displayed in Supplementary Table 5. One example is a locus on chromosome 6 which colocalizes with asthma (6_90220794_T_C) and Crohn’s disease (6_90263440_C_A) suggesting that the two diseases may share common genetic etiology at this locus.

To demonstrate the value of colocalization evidence, we examined coding variants that were fine-mapped to single-variant resolution, and which colocalized with a molecular QTL for the same gene (729 variants, Supplementary Table 6). Such cis-variants make good genetic instruments for testing the causal effect of the molecular phenotype on disease ^28^, and the ratio of coefficients for the cis-variants is an estimate of the effect size of the molecular phenotype on disease. Using this approach we identified several known gene-trait associations. For example, missense variant rs34324219 is causal of changes in *TCN1* RNA and protein expression in whole blood ^15,17^ and also colocalizes (H4>0.99) with pernicious anemia, a disorder in which too few red blood cells are produced due to vitamin B12 deficiency. *TCN1* encodes the protein haptocorrin (also known as Transcobalamin-1) which binds vitamin B12 and is involved in its uptake ^29^. Also, splice region variant rs1893592 causes increased expression of *UBASH3A* in most GTEx tissues, including thyroid. This signal colocalizes (H4>0.87) with self-reported treatment using the thyroid hormone sodium levothyroxine. Hypothyroidism is a common comorbidity with type 1 diabetes, for which there is strong evidence that *UBASH3A* is causal ^30^. Finally, the synonymous variant rs2228079 is the only credibly causal variant for an eQTL associated with altered *ADORA1* expression in whole blood (eQTLGen) and colocalizes with asthma in UK Biobank (H4>0.99). *ADORA1* encodes a type of adenosine receptor, a class of proteins targeted by the approved drug (Theophylline) for the treatment of asthma.

Colocalization also provided strong genetic evidence for some less well known gene-disease associations (Supplementary Table 7). One example is splice region variant rs11589479, which causes increase in *ADAM15* expression in several monocytes states and also colocalizes (H4=0.99) with Crohn’s disease ^31^. *ADAM15*, a disintegrin and metalloproteinase, is strongly upregulated in colon tissues from inflammatory bowel disease patients compared to healthy controls and plays a role in leukocyte trans-migration across epithelial and endothelial barriers as well as the differentiation of regenerative colonic mucosa ^32^.

### A machine learning model prioritizes genes at gold-standard loci

We next developed a “locus to gene” model (L2G) to prioritize causal protein-coding genes at GWAS loci by integrating our catalog of fine mapping associations with relevant functional genomics features. We first manually curated a set of 445 gold standard positive (GSP) genes at GWAS loci for which we are confident of the causal gene assignment (Supplementary Table 8, see methods). The selected genes are based on (i) expert domain knowledge of strong orthogonal evidence or biological plausibility; (ii) known drug target-disease pairs; (iii) experimental alteration from literature reports (e.g. nucleotide editing); (iv) observational functional data (e.g. colocalizing molecular QTLs, colocalizing epigenetics marks, reporter assays) (Supplementary Table 9). Next, we defined locus-level predictive features from four evidence categories: in silico pathogenicity prediction from VEP and PolyPhen, colocalization of molecular QTLs, gene distance to credible set variants weighted by their fine-mapping probabilities, and chromatin interaction (Supplementary Table 10). The chromatin interaction data comprised promoter-capture Hi-C from 27 cell types ^33^, FANTOM enhancer-TSS pairwise cap analysis of gene expression correlation^34^; and DNase I hypersensitive site-gene promoter correlation^35^. Then, using a nested cross-validation strategy, we trained a gradient boosting model to distinguish GSP genes from other genes within 500 kb at the same loci (see methods).

The L2G model produced a well calibrated score, ranging from 0 to 1, which reflects the approximate fraction of GSP genes among all genes above a given threshold (Figure 2). At a classification threshold of ≥0.5, the full model correctly identified 238 out of 445 true positives with 86 false positives (average precision = 0.65; Table 1). We compared the full model against a naive nearest gene classifier (closest gene footprint and closest TSS), which selects the closest gene to each lead variant, and thus does not make use of other candidate variants from fine-mapping. The naive nearest gene classifier identified more true positives at the same threshold (268 out of 445) but at the cost of identifying 2.4 times more false positives (207) (Average precision=0.37). Hence the full L2G model has higher precision with a small reduction in recall.

**Figure 2:**
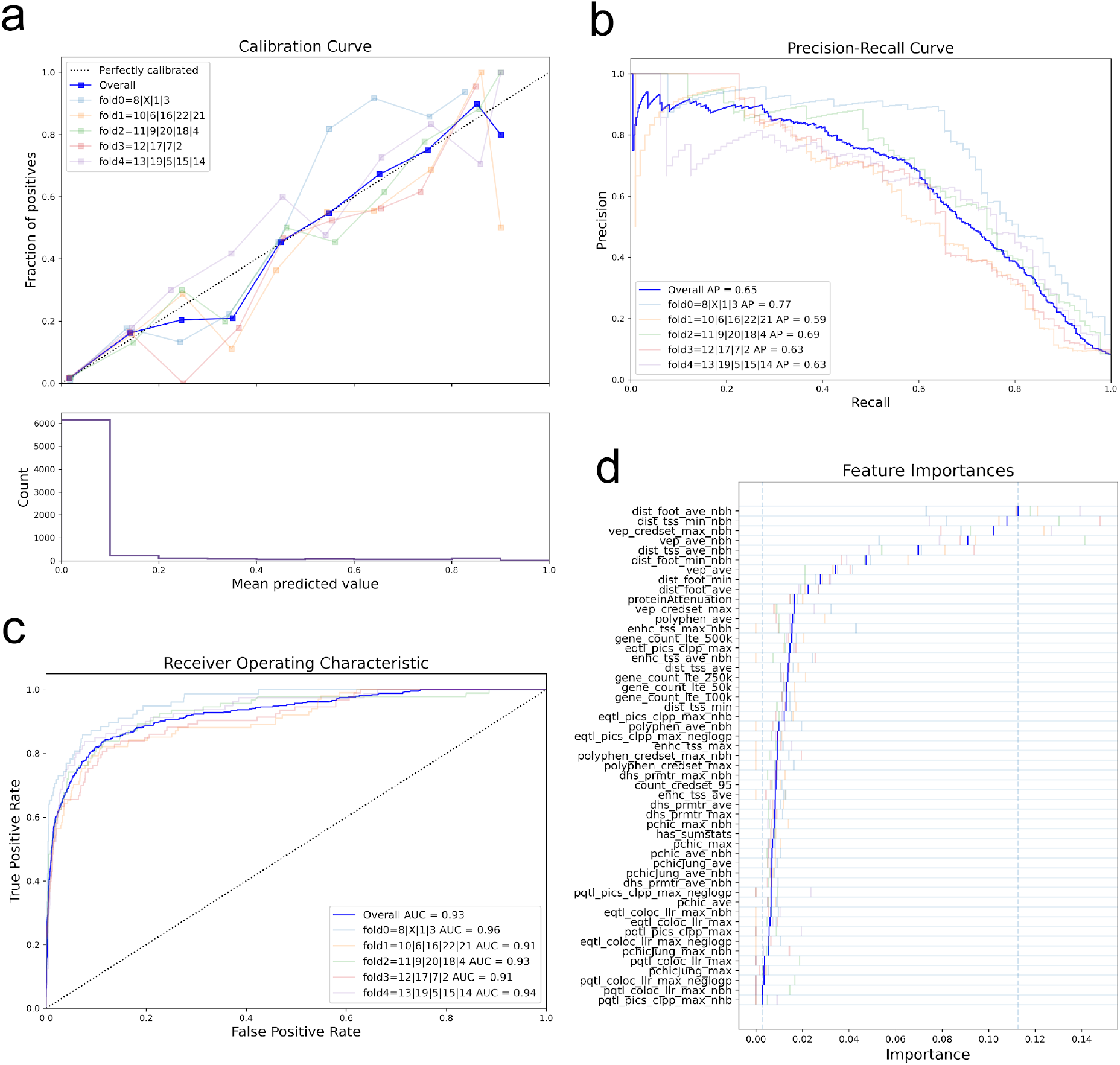
Performance of the locus-to-gene (L2G) model. (a) Calibration curve, showing (top) the fraction of all GSP genes found as positives at different L2G score thresholds (mean predicted value), and (bottom) the count of genes in each L2G score bin. (b) The precision-recall curve and (c) the receiver-operator characteristic curve for identifying GSP genes from among those within 500 kb at each locus. (d) The *Relative Importance* of each predictor in the L2G model.

To identify which features are most important in predicting GSP genes, we retrained the model to include features from only one of the four evidence categories at a time (leave-one-group-in analysis). No individual feature set gets a higher ‘Average Prediction’ score as the full model (Table 1). Our ‘mean distance’ feature which aggregates across all the variants in the credible set and weighs by their posterior probability was the most predictive (average precision=0.62) followed by *in silico* pathogenicity prediction evidence (average precision=0.48), molecular QTL colocalization (average precision=0.36) and chromatin interaction (average precision=0.26) (Table 1, Leave-one-group-in section). Note that the ‘mean distance’ feature is distinct from a ‘naive closest gene distance’ feature because of the weighting across a credible set to the most likely SNPs, and thus manages to discard many false positives (FP_*mean distance*_ = 98 vs FP_*naive closest footprint gene*_ = 207 and FP_*naive closest TSS gene*_ = 195). Within the mean distance features tested, whether the gene was the closest at the locus using a gene footprint distance metric averaged over the credible set and whether the gene was the closest at the locus using the minimum gene-TSS distance over the 95% credible set, had the highest relative feature importances (Figure 2d). Thus, when using distance as a predictor of causal genes, the distance relative to other genes is more important than the absolute distance.

**Table 1.** Classification performance for feature groups. Performance characteristics of the full model are shown at the top, and analyses for individual groups of features are shown in sections below. Counts are shown for true positives (TP), false positives (FP), true negatives (TN), and false negatives (FN). * Mean Distance aggregates across all the variants in the credible set and weighs by their posterior probability.

| Features | Average<br>precision | AUC | Precision | Recall | TP | FP | TN | FN | Sensitivit<br>y | Specificit<br>y | FDR | GSP<br>count | GSN<br>count |
| --- | --- | --- | --- | --- | --- | --- | --- | --- | --- | --- | --- | --- | --- |
| Full model | 0.65 | 0.93 | 0.73 | 0.53 | 236 | 86 | 6429 | 209 | 0.53 | 0.99 | 0.27 | 445 | 6515 |
| **Naïve closest gene classification** |  |  |  |  |  |  |  |  |  |  |  |  |  |
| Closest footprint | 0.37 | 0.79 | 0.56 | 0.6 | 268 | 207 | 6308 | 177 | 0.6 | 0.97 | 0.44 | 445 | 6515 |
| Closest TSS | 0.34 | 0.76 | 0.56 | 0.55 | 246 | 195 | 6320 | 199 | 0.55 | 0.97 | 0.44 | 445 | 6515 |
| **Leave-one-group-in** |  |  |  |  |  |  |  |  |  |  |  |  |  |
| Mean Distance* | 0.62 | 0.91 | 0.69 | 0.49 | 219 | 98 | 6417 | 226 | 0.49 | 0.98 | 0.31 | 445 | 6515 |
| Interaction | 0.26 | 0.79 | 0.55 | 0.05 | 23 | 19 | 6496 | 422 | 0.05 | 1 | 0.45 | 445 | 6515 |
| Molecular QTL | 0.36 | 0.85 | 0.62 | 0.18 | 79 | 49 | 6466 | 366 | 0.18 | 0.99 | 0.38 | 445 | 6515 |
| Pathogenicity<br>prediction | 0.48 | 0.76 | 0.7 | 0.43 | 191 | 80 | 6435 | 254 | 0.43 | 0.99 | 0.3 | 445 | 6515 |
| **Leave-one-group-out** |  |  |  |  |  |  |  |  |  |  |  |  |  |
| Mean Distance* | 0.47 | 0.77 | 0.69 | 0.43 | 191 | 84 | 6431 | 254 | 0.43 | 0.99 | 0.31 | 445 | 6515 |
| Interaction | 0.65 | 0.93 | 0.73 | 0.53 | 234 | 85 | 6430 | 211 | 0.53 | 0.99 | 0.27 | 445 | 6515 |
| Molecular QTL | 0.65 | 0.93 | 0.74 | 0.54 | 239 | 86 | 6429 | 206 | 0.54 | 0.99 | 0.26 | 445 | 6515 |
| Pathogenicity<br>prediction | 0.63 | 0.92 | 0.71 | 0.5 | 222 | 91 | 6424 | 223 | 0.5 | 0.99 | 0.29 | 445 | 6515 |

We also assessed the unique contribution of each evidence type by leaving out one group of features at a time. Consistent with the leave-one-group-in analysis, dropping our mean distance features had the largest impact on prediction (*average precision* change from 0.65 to 0.47), followed by *in silico* pathogenicity prediction (*average precision* down to 0.63) (Table 1). Notably, when molecular QTL colocalization evidence was removed from the model we saw similar classification results, with 3 fewer true positives identified, and no net change in the Gold Standard Negatives (GSN)(Supplementary Table 11a). There are various possible reasons for this: the colocalization score may be redundant with some of our other features; we may lack the relevant tissue- or context-specific QTLs; or we may have obscured the utility of colocalization information by using a cross-tissue colocalization score. We also used a measure of *continuous reclassification improvement* to evaluate prediction changes across all possible classification thresholds. Here, adding molecular QTL colocalization evidence resulted in a net 4.7% GSPs having an increased prediction score and a net 42.2% GSNs having a decreased score (Supplementary Table 11b). This suggests that whilst our colocalization features do not provide sufficient evidence to support novel positives, lack of colocalization accurately identifies negative gene assignments. Removing chromatin interaction features resulted in a minor reduction in model performance (net 2 fewer GSPs) (Table 1).

The low predictiveness of features apart from distance relates in part to their lower genome coverage. For distance features, most sentinel variants have at least 1 gene within 500 kb, but for pathogenicity, molecular QTL colocalization and chromatin interaction, coverage of variants was low (Supplementary Figure 2). Only a small proportion of studies had summary statistics available, limiting our ability to use *coloc* to perform a colocalization analysis (only 3% of all loci had *coloc* derived evidence). Our complimentary colocalization method, using a reference LD-panel to approximate summary statistics (the PICS method), increased the total number of loci with colocalization evidence to 19%. Evidence from pQTLs was very sparse at <1% coverage, which may account for its very low feature importance (Supplementary Figure 2).

### Gene prioritization across all trait-associated loci

We used the trained L2G model to prioritize causal genes across all 133,441 trait-associated GWAS loci in our repository. At a classification threshold of 0.5, 55.4% (n=74,096) of all loci had a single gene prioritized whereas only 1.4% (n=1,907) had 2 or more genes prioritized (Supplementary Figure 3). 43.2% of loci did not reach the classification threshold. Across all diseases, genes prioritized by the model were 7.8 times more likely (95% CI: [6.5, 9.3]) to be supported by literature evidence identified by text mining (Supplementary Table 12). Genes prioritized by the naive classifier using the closest gene footprint from the sentinel variant were also enriched (5.6 times, 95% CI: [4.7, 6.6]) but not as highly as the full model (p-value=0.008 against null-hypothesis logOR_Full model_ = logOR_Naive mode_, Welch t-test).

In order to benchmark the L2G versus the distance based classifier, we tested whether prioritized gene-diseases were enriched for known drug target-indication pairs across different clinical phases according to the ChEMBL database. Genes prioritized by the model were enriched with OR 7.4, 8.5 and 8.1 (95% CI: [5.7, 9.4], [6.3, 11.3], [5.7, 11.5]) across clinical trial phases ≥2, ≥3 and 4, respectively (Supplementary Table 13). Using a naive classifier we saw lower odds ratio point estimates but with overlapping confidence intervals (OR 5.3 [4.2, 6.7], 6.4 [4.8, 8.5] and 6.7 [4.8, 9.3]) (Supplementary Figure 4). Thus the prioritisation using the L2G model both recapitulates the established enrichment of GWAS loci for known drugs^11^ but also demonstrates that fine-mapping and colocalization combined with the L2G approach improves on their approach, and hence is likely to also improve success in identifying novel drug targets.

## Discussion

To address the challenges of translating GWAS signals to biological insights, we developed a pipeline to format, harmonize, and aggregate human trait and disease GWAS, molecular QTLs and functional genomics data in a consistent way, providing statistical evidence for target prioritization across the entirety of GWAS traits and diseases. We then trained a machine learning model that integrates fine-mapping and functional genomics data to prioritize likely causal variants and genes at 133,441 trait-lead variant disease associations. The L2G score output by the model represents the likelihood that a gene is causal for that trait, subject to the limitations of our gold standard positive training data, and thus allows genes at all trait-associated loci to be ranked by the relative strength of their evidence. Under cross-validation, the model resulted in a 58% reduction in the number of false-positives detected (improved precision), at the cost of missing 11% of the gold-standard positives (reduction in recall). The top genes prioritized by the L2G score recover known relationships, including disease-gene pairs with approved drugs, as well as novel disease-drug target associations that suggest potential novel therapeutic targets to pursue.

The strength of our machine learning approach stems from the systematic application of fine-mapping to obtain per-variant probabilities prior to gene assignment. Sentinel variants discovered by GWAS may not be the causal variant ^36^; by aggregating functional data across the credible set we incorporate information from all plausible causal variants at the locus. Using a supervised learning method allowed us to efficiently combine heterogeneous functional datasets into a single model. The L2G score output by our model is well calibrated, meaning that it can be interpreted as a probability and thus the evidence supporting a gene assignment can be compared both within and between loci.

A limitation of our approach is that it requires a large number of high-quality gold standards to train the model, and each source of gold standards will have biases. For example, when we compared the dataset of drug targets from CHEMBL retrospectively mapped to GWAS loci to the manually curated datasets (mainly focused on the closest genes and those with known missense variants), we found that distance and VEP features performed much better in the manually curated datasets (Supplementary Figure 5), emphasizing the need to curate less-biased datasets. Using varied sources may help mitigate some source-specific biases, but manually curated allele-gene pairs are intrinsically more likely to be close to each other. Future gold-standard training data should represent a range of possible molecular mechanisms. The reliance on large amounts of training data influenced the design of our model. To avoid stratifying gold-standards into smaller subgroups, we trained the model across all diseases at once and using functional data ascertained from different tissues/cell types aggregated into a single feature. This means that the model is not currently able to specifically leverage the tissues/cell types that are most relevant for a given disease.

The outputs of our analyses can be viewed in the Open Targets Genetics portal (https://genetics.opentargets.org), a user-friendly web interface that supports visualisation of fine-mapping and L2G scores for individual variants and genes across 133,441 trait-lead variant GWAS associations. The portal also offers other features including disease-disease and disease-molecular traits colocalization analyses across ~3,600 GWAS summary statistics and 92 tissue and cell type-specific molecular QTL summary statistics to identify traits and diseases that share common genetic susceptibility mechanisms.The portal will regularly be updated with new GWAS summary statistics both from Europeans and non-European ancestries as well as QTLs and functional genomic data from a wider range of tissues and cell types. Planned enhancements include displaying tissue- and cell type-specific enrichments for each included trait, using methods such as CHEERS ^37^ that leverage functional annotations. These enrichments will also be used to improve the L2G model by using functional genomics data from tissues that are most relevant to each disease and trait. Our repository of gold-standard gene assignments will be expanded as more evidence arises. In particular, we encourage scientists from the genetics community to contribute to this repository, since having diverse evidence sources can partially address the bias that comes with manually curated sets.

## Methods

### Summary statistics based fine mapping

We harmonised summary statistics to ensure alleles and effect directions were consistent across studies, and removed variants with low confidence estimates (minor allele count < 10). We identified independently associated loci for each study using Genome-wide Complex Trait Analysis Conditional and Joint Analysis (GCTA-COJO; v1.91.3) ^21^. UK Biobank genotypes down-sampled to 10k individuals were used as a linkage-disequilibrium (LD) reference for conditional analysis ^38^. We considered a locus to be independently associated if both marginal and conditional p-values were less than 5e^−8^. For each independent locus, we produced a set of summary statistics that are conditional on all other independent loci ±2Mb from the sentinel variant. Using the conditional set of summary statistics, we computed approximate Bayes factors ^39^ from the beta and standard error for each SNP, with a variance prior (W) of 0.15 for quantitative traits and 0.2 for binary traits, and determined variant posterior probabilities (PP) assuming a single causal variant as: PP = SNP BF / sum(all SNP BFs) for all SNPs within a ±500Kb window. We considered any variant with a PP > 0.1% as being in the credible set.

### Linkage-disequilibrium based fine mapping

In addition to the above fine mapping analysis, we conducted a complementary LD based approach which allowed us to leverage information from studies that lack full summary statistics. For each independent locus, we identified all variants in LD with the sentinel variant (R^2^>0.5 in ±500Kb window). LD was calculated in 1000 Genomes phase 3 data ^40^ by mapping the GWAS study ancestries to the closest super population ^41^, taking a sample size weighted-mean of the Fisher Z-transformed correlations in the case of multi-ancestry studies. We then used the Probabilistic Identification of Causal SNPs (PICS) method to estimate the PP that each variant is causal based on the LD structure at each locus ^23^. As above, we kept all variants with PP > 0.1%.

### Colocalization analysis

Molecular QTL summary statistics were acquired from the EBI eQTL Catalogue ^16^, GTEx (v7) ^14^, eQTLGen ^15^ and Sun et al. protein QTLs ^17^. Summary statistics were restricted to be ±1Mb from the gene transcription start site (TSS). We pre-processed and fine mapped molecular QTL summary statistics using the same method described above for GWAS studies. However, we used less stringent criteria for the inclusion of QTL lead variants, requiring minor allele count ≥ 5 and adjusted for multiple testing using a Bonferroni correction of p < 0.05 / number of variants tested per gene.

For GWAS studies with summary statistics, we performed a colocalization analysis if there was at least 1 variant overlapping between the GWAS and molecular trait 95% credible sets (prior evidence for colocalization). We conducted colocalization of summary statistics using the coloc package (v.3.2-1) ^27^ with default priors. Given that there is prior evidence for colocalization, these parameters will give conservative estimates. As with the fine mapping pipeline, we used summary statistics conditional on all other independent loci within ±2Mb and restricted the coloc analysis to a ±500Kb window around each sentinel variant. A minimum of 250 intersecting variants were required for analysis.

For GWAS studies without summary statistics, we performed an alternative colocalization analysis using the LD-based PICS fine mapping sets. Colocalization was approximated by taking variants that intersect at pairs of GWAS and molecular trait loci, and summing the product of the PPs.

### Pre-processing of functional genomics data for L2G prioritization

We used 4 main classes of evidence to prioritize genes: (i) variant pathogenicity in silico predictions; (ii) colocalization with molecular trait quantitative trait loci (QTL); (iii) chromatin conformation; (iv) linear genomic distance from variant to gene.

We used *in silico* pathogenicity predictions to estimate the effect of variants on gene transcripts and protein function. Firstly, we incorporated Variant Effect Predictor (VEP) ^24^ transcript consequences. We mapped VEP’s impact ratings of High, Moderate, Low to scores of 1.0, 0.66, 0.3 (respectively), and included an additional four consequences (intronic, 5’ UTR, 3’ UTR, nonsense-mediated mRNA decay transcript variants) with a score of 0.1 as we expected them to have predictive value through their functional consequences on mRNA transcription, secondary structure and translation. For each variant-gene pair we took the maximum score across transcripts. In addition to VEP we included PolyPhen-2 pathogenicity scores representing the probability that a non-synonymous substitution is damaging ^42^.

Chromatin interaction data were from promoter-capture Hi-C, FANTOM enhancer-TSS correlation, and DNase-hypersensitivity enhancer-promoter correlation. Each of the data points in these datasets is represented as a pair of interacting genomic intervals and an association statistic. We retained interval pairs with one end encompassing an Ensembl gene Transcription Start Site (TSS)^43^ and the other end containing any variant in Gnomad 2.1 ^44^, resulting in variant-gene pairs with a dataset-specific association statistic.

We included two genomic distance metrics as it has been shown that, despite notable contrary exceptions, linear distance is a good predictor of candidate causal genes ^45^. First, the distance from each variant to all gene TSSs is included. Second, the distance from each variant to each gene’s footprint, where the footprint is any position between the start and end positions of the gene. For both metrics the canonical transcript is used, as defined by Ensembl for protein-coding genes within a ±500Kb window around each variant.

### Derivation of locus-to-gene prioritization features

We next combined our fine mapping and functional genomics data to create features to prioritize candidate causal genes at each trait-associated locus (locus-to-gene scoring) (Supplementary Table 10).

Except for molecular trait colocalization evidence, each functional genomics dataset is variant-centric, meaning they give variant-to-gene scores. We convert variant-centric scores into locus-to-gene scores by aggregating over credible variants identified through fine mapping. For GWAS studies with summary statistics available we used ABF credible sets, otherwise we used LD-based PICS credible sets. We implemented two complementary methods for aggregating over credible sets. Firstly, we took a weighted sum of scores across all variants identified by fine mapping (PP > 0.01%) using PP of causality as weights (Equation 1). Secondly, we took the maximum score for any variant in the 95% credible set (Equation 2).

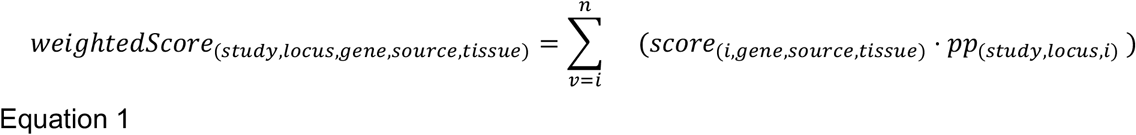

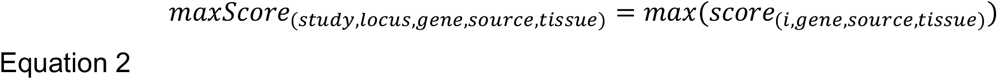

Molecular trait colocalization evidence is a locus-centric score. We included both summary statistic derived *coloc* evidence (Equation 3) and LD-derived colocalization evidence as features. Each GWAS signal may have colocalization estimates from multiple independent molecular trait signals (each conditional on the others), we therefore took the maximum score across estimates. Given that evidence against colocalization (*h*_*3*_) cannot be directly estimated without full summary statistics, this term was dropped for the LD-derived colocalization feature (Equation 4).

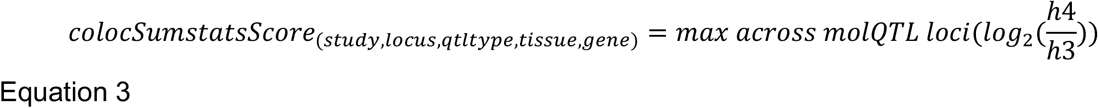

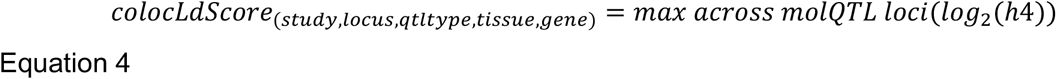

For functional genomics datasets with measurements in multiple tissues (or cell types), we calculated the locus-level feature for each tissue separately and took the maximum across tissues (Equation 5).

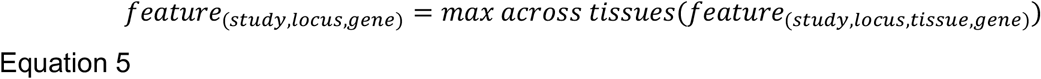

We next wanted to provide the model with information about other genes at each locus (termed the *neighbourhood* feature). This allows the model to learn whether a given gene has, for example, the highest colocalization score compared to others at the locus. To do this we divided each feature by the maximum score across genes at that locus (Equation 6).

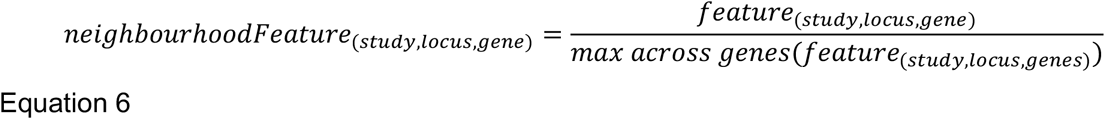

### Curation of a GWAS gold-standard training dataset

We next assembled a repository of published GWAS loci (https://github.com/opentargets/genetics-gold-standards) for which we have high confidence that the gene mediating the association is known. Gold-standard evidence were grouped into 4 classes: (i) *expert curated* loci with strong orthogonal evidence or biological plausibility; (ii) *drug* loci inferred from known drug target-disease pairs; (iii) loci inferred from *experimental* alteration (e.g. nucleotide editing); (iv) loci inferred from *observational* functional data (e.g. colocalizing molecular QTLs). We also assigned each gold-standard a confidence rating of *high*, *medium* or *low* depending on our assessment of the strength of supporting evidence.

We started by compiling existing gold-standard examples from the literature. 227 curated metabolite QTLs were sourced from Stacey *et al ^45^* and a further 136 loci were curated by Eric Fauman with strong biological plausibility (Supplementary Table 6). We then ascertained 57 genes with “causal” or “strong” *observational* data from the Type 2 Diabetes Knowledge Portal Effector Genes table, this equates to genes with: a confirmed causal coding variant; or at least two of the following: (i) a likely causal coding variant, (ii) >1 piece of regulatory evidence, >1 piece of perturbation evidence ^46^. We added a further 48 disease-causal genes curated from the literature. These were mainly GWAS associated loci that were fine-mapped and colocalized with eQTL and epigenomic features in disease-relevant tissues in order to prioritize likely functional variants and their causal genes. These results were then functionally validated using experiments such as reporter assays and CRISPR/Cas9 genome editing.

In addition to literature sourced loci, gold-standard evidence was generated based on known drug-target-indication associations curated in ChEMBL in clinical trial phase II, III or IV ^47^. Drugs that bind a protein complex, rather than a single protein, were removed unless the binding subunit was known.The ChEMBL evidence was combined with the genetics features to identify loci with known drug targets. Gold-standards derived from phase II, III and IV drug targets were assigned a confidence of *low*, *medium* and *high*, respectively. Additionally, confidences were adjusted to indicate the distance of the sentinel variant to the drug target, variant-gene distances of < 500, 250, 100Kb kb were assigned confidences *low*, *medium* and *high*, respectively.

Duplications were removed from the Gold-standard positives (GSP) list so that GWAS allele-gene pairs never occurred more than once in the training data. The same gene could occur as a GSP more than once if the associated alleles were independent, i.e. if no variants overlapped between their credible sets (using all variants with PP > 0.1%). All non-GSP genes in the training data at the locus (±500kb) were set as gold-standard negatives (GSN). GSNs genes were subsequently removed if they had a stringDB score ≥ 0.7 with the GSP at the same locus, the aim being to remove alternative explanations for the association between trait-associated allele and gene. This resulted in a total of 229 GSNs being removed (out of a total of 9,171). A total of 445 GSP were included in the final training data.

### Supervised learning of locus-to-gene features

We used all GWAS loci with high or medium confidence gold-standard evidence (445 loci) to train an XGBoost gradient boosting classifier ^48^ using a binary logistic learning objective function. Nested cross-validation (CV) as implemented in scikit-learn was used to maintain independence of the training and test data and to tune hyperparameters. The outer CV consisted of 5 folds split by chromosomes so that each group contained an approximately equal number of GSPs. Within each fold, we used a random parameter search to train 1000 models, which were assessed using a *balanced accuracy* metric averaged over 5 randomly split inner folds.

For each group of features included in the main model, we conducted sub-analyzes whereby either only that feature group was included (leave-one-group-in), or everything except that feature group was included (leave-one-group-out). This allowed us to evaluate the relative performance of each feature group individually. Additionally, we output the *Relative Importance* of each feature as implemented in the XGBoost model ^49^.

### Model internal validation

Our cross-validation approach produces separate models for each of the 5 outer folds. We evaluated the performance of each model against the remaining 20% of loci not used for training. We used *average precision* and *area under the receiver operator curve (AUC)* metrics to assess the classification across the full range of prediction probabilities outputted by the model. We also assess the performance of the model after applying a hard threshold of >0.5 (>50% confidence that the characteristics of the observed locus is consistent with being a gold-standard positive locus).

We compared the relative performance of leave-one-group-in and leave-one-group-out models by calculating the *net reclassification improvement* (NRI) of loci compared to the full model ^50^. NRI measures the number of GSP loci that move above the classification threshold (>0.5), compared to GSN that move below, when the model is updated. We also calculate *continuous NRI (cNRI)*, the sum of the percentage of GSPs with classification scores that move in the correct direction vs. GSNs that move in the wrong direction (towards higher scores) ^51^.

### Model external validation with literature evidence

We benchmarked the L2G assignment against independent gene-disease associations scored by literature mining in the Open Targets Platform. We excluded any publications for studies curated in GWAS Catalog to ensure independence of the training data. We restricted analyses to a subset of 22 prioritized diseases (Coronary artery disease, Breast carcinoma, Prostate carcinoma, Acute lymphoblastic leukemia, Inflammatory bowel disease, Crohn's disease, Ulcerative colitis, Rheumatoid arthritis, Osteoarthritis, Type I diabetes mellitus, Hypothyroidism, Psoriasis, Atopic eczema, Asthma, Alzheimer's disease, Parkinson's disease, Ankylosing spondylitis, Celiac disease, Gout, Multiple sclerosis, Systemic lupus erythematosus). For each disease, we constructed a 2×2 contingency table of ‘gene prioritised by L2G model (score > 0.5)’ and ‘gene prioritised by Open Targets literature evidence (top decile [>0.52])’. Only genes scored by the L2G model (±500kb of a sentinel GWAS variant) were included in the contingency table. We calculated enrichment and statistical significance using Fisher’s exact test.

### Enrichment of known drug targets

We calculated drug target enrichment using known target-indication pairs curated in ChEMBL (accessed: 2019-03-25). We constructed a single 2×2 contingency table pooling across all indications, which consisted of ‘gene prioritized by L2G model (score > 0.5)’ and ‘gene is known target of drug for indication matched to GWAS disease phenotype’. GWAS studies were only included if they could be mapped to a ChEMBL indication (matched using Experimental Factor Ontology) and that indication has a known drug that can be mapped to a protein-coding gene that was scored by the L2G model. Enrichment was calculated by Fisher’s exact test.

## Data availability

Our results are freely available through a web portal (genetics.opentargets.org), GraphQL API or through bulk download. GWAS gold standard genes: github.com/opentargets/genetics-gold-standards.

## Acknowledgements

The authors would like to thank Ellen McDonagh, Joe Maranville, and David Hulcoop for their useful feedback to improve the paper, Helen Parkinson, and Jacqueline MacArthur for their support with the GWAS Catalog data. This research has been conducted using the UK Biobank Resource. This work was funded by Open Targets. EM work was funded by JDRF (4-SRA-2017-473-A-N) to the Diabetes and Inflammation Laboratory, University of Oxford.

## Author contributions

MG, JS, EM, ID wrote the manuscript. EM conducted the analysis and designed and built the ML model. EM, EMS, MG prioritised GWAS studies for curation by GWAS Catalog. EM, MC, AB, JH, EP curated and processed the GWAS and functional genomics data, EF, EM, MG curated the gold standards. GP, AM, LF, AH, EP designed and implemented visualisations for analysis. DO performed additional analysis. ID, MG, JAT, JCB conceived and supervised the study. MAK generated Figure 1. MG, EM, EMS, DW, EP worked on the biological questions and the underlying visualisations in the portal.

