## Supplementary Figures for "Open Targets Genetics: An open approach to systematically prioritize causal variants and genes at all published human GWAS trait-associated loci" (from another version of this preprint: version 1, 2020-09-17)

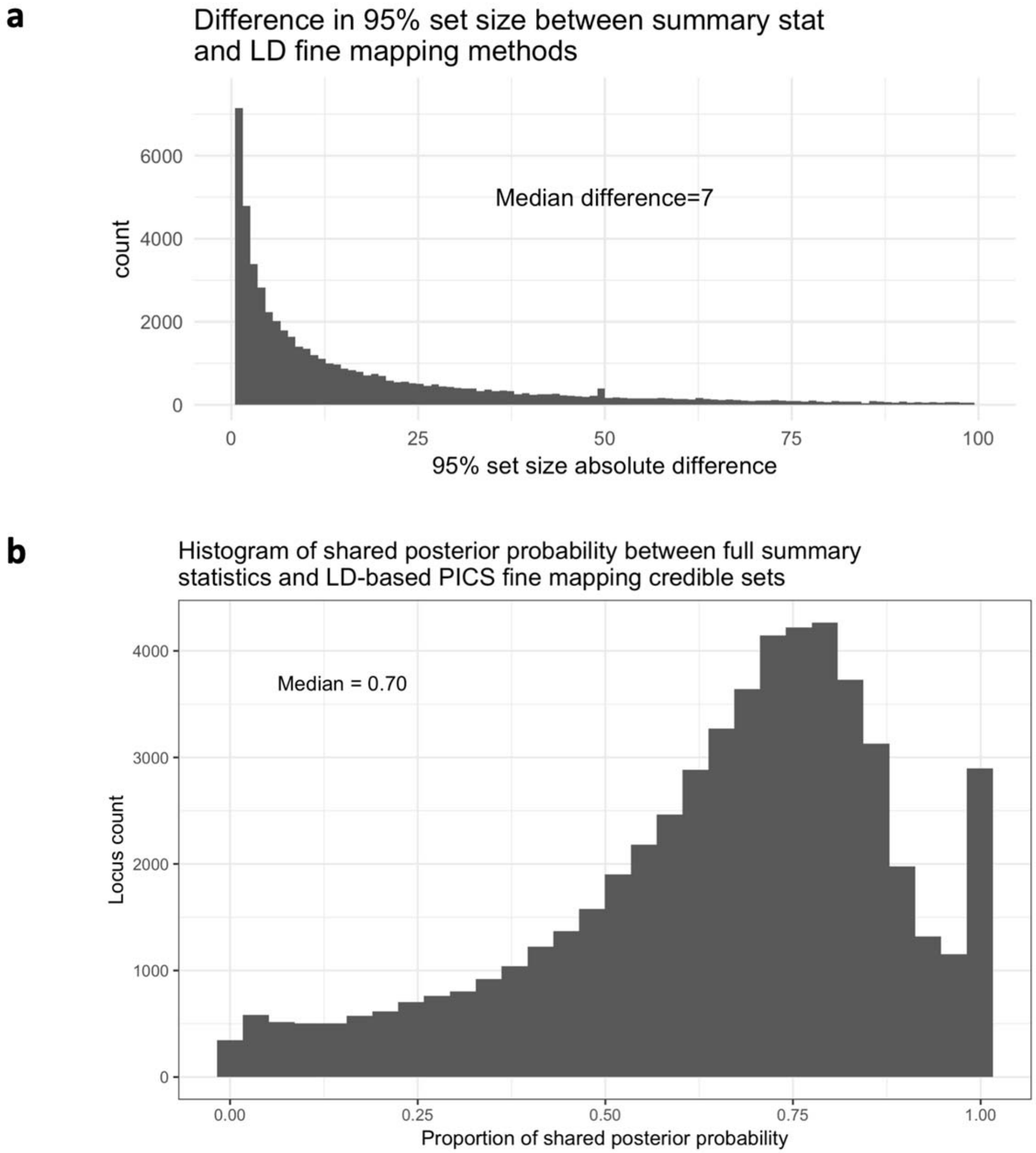


**Supplementary Figure 1**: Difference between fine-mapping methods. (a) Histogram of the absolute difference in the number of variants in the 95% credible set across all loci. The median absolute difference was 7 variants. (b) Histogram of the posterior probability at a given locus that is contained in variants shared between the 95% credible sets of the two methods, determined for all loci. The median shared variant probability was 0.70.

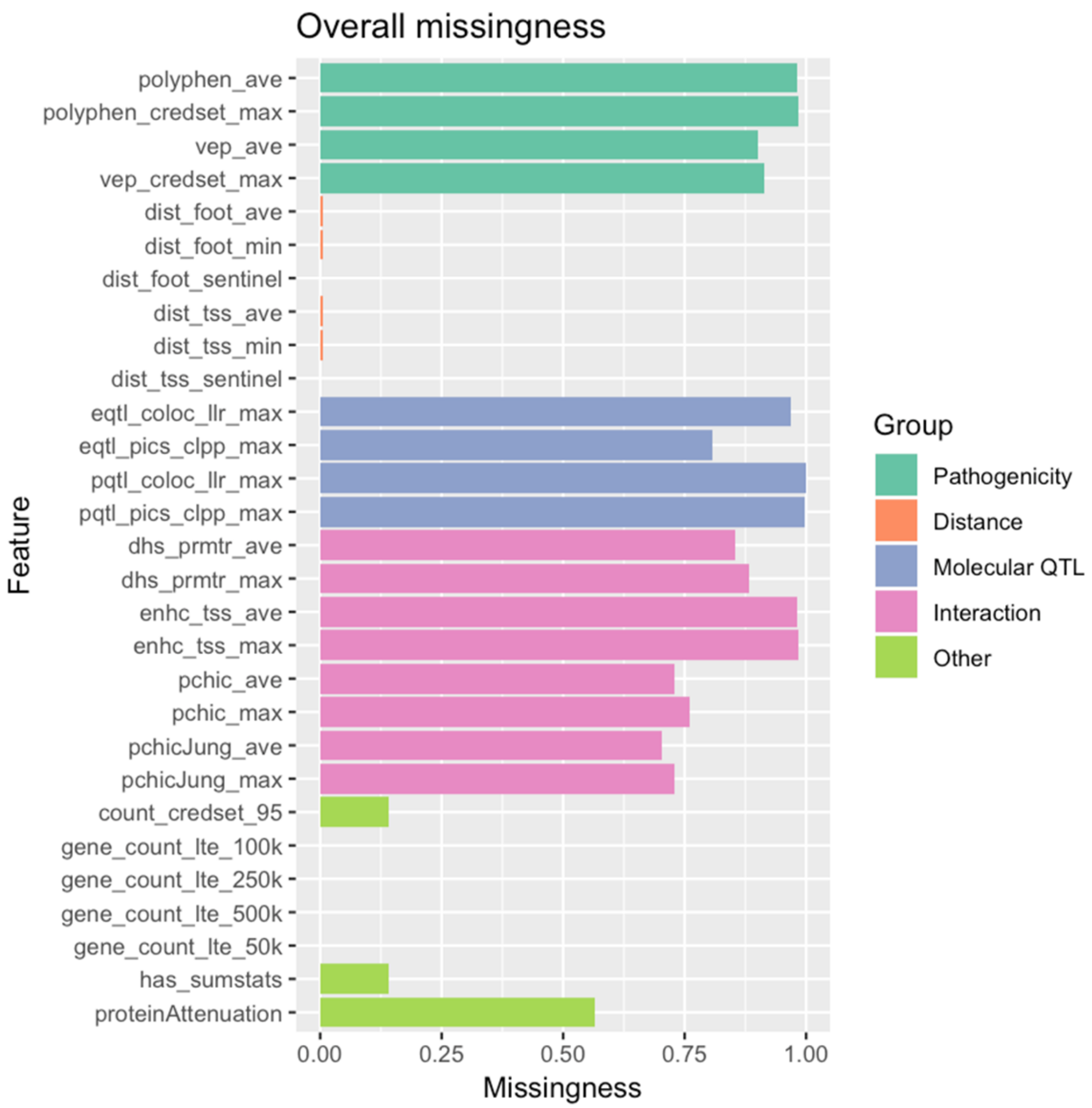


**Supplementary Figure 2**: Missingness. The fraction of variants with missing values (no annotation in that category) is shown for representative input features of the L2G model.

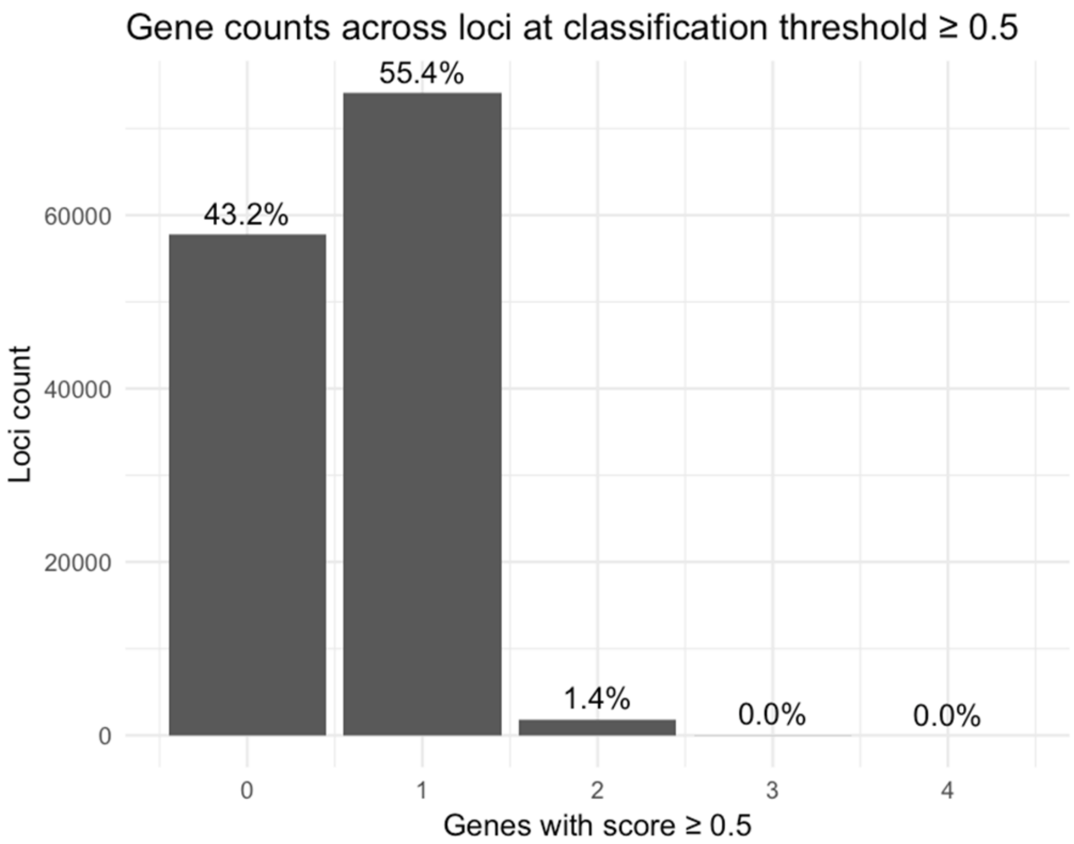


**Supplementary Figure 3**: Histogram of the number of prioritized genes (having L2G score >= 0.5) at each locus. Very few loci have more than one gene prioritized.

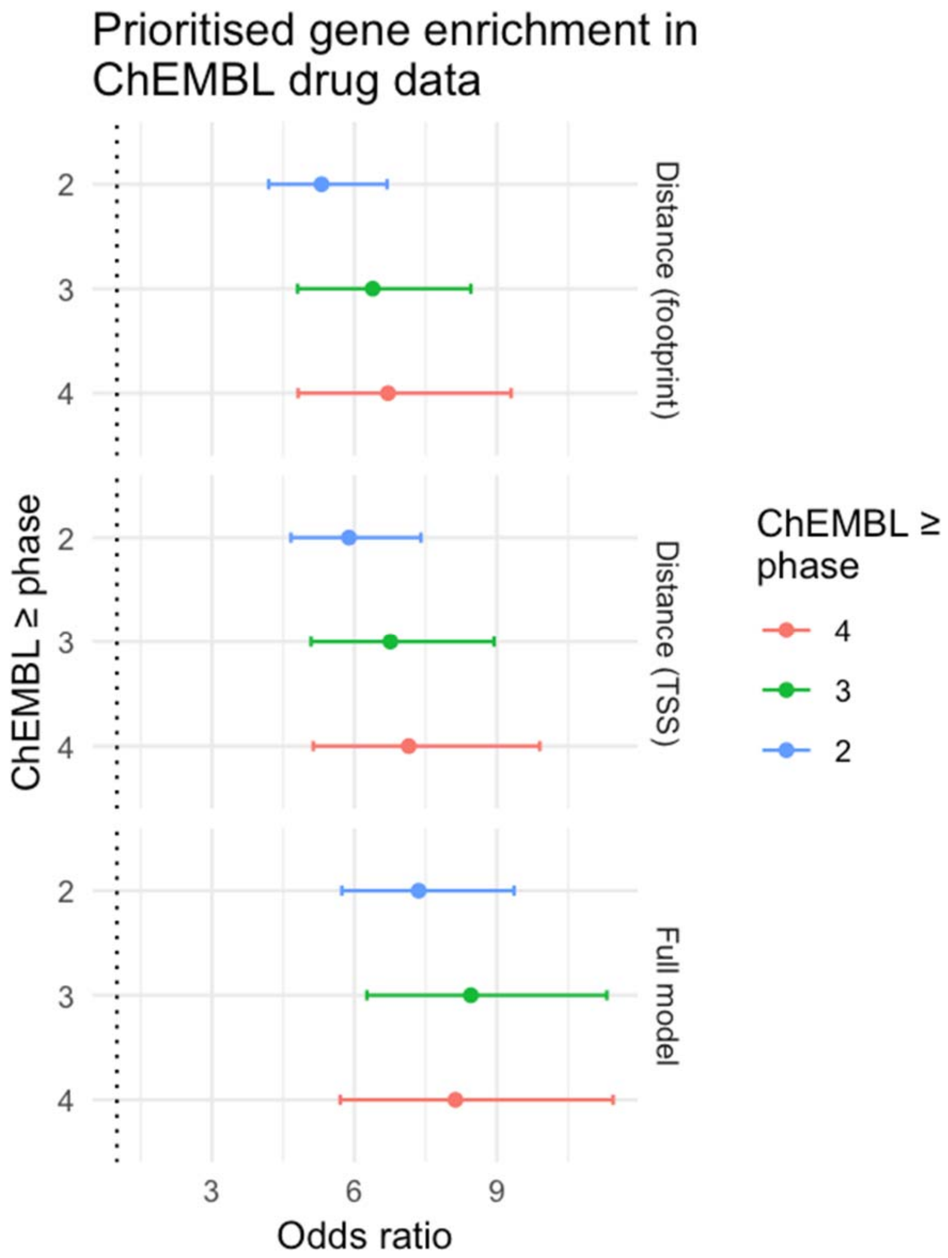


**Supplementary Figure 4**: Enrichment of genes with model score >= 0.5 for the distance-only models (top 2) or the full L2G model (bottom), stratified by whether the gene is a known drug target in ChEMBL phase >= 2, >= 3, or >= 4.

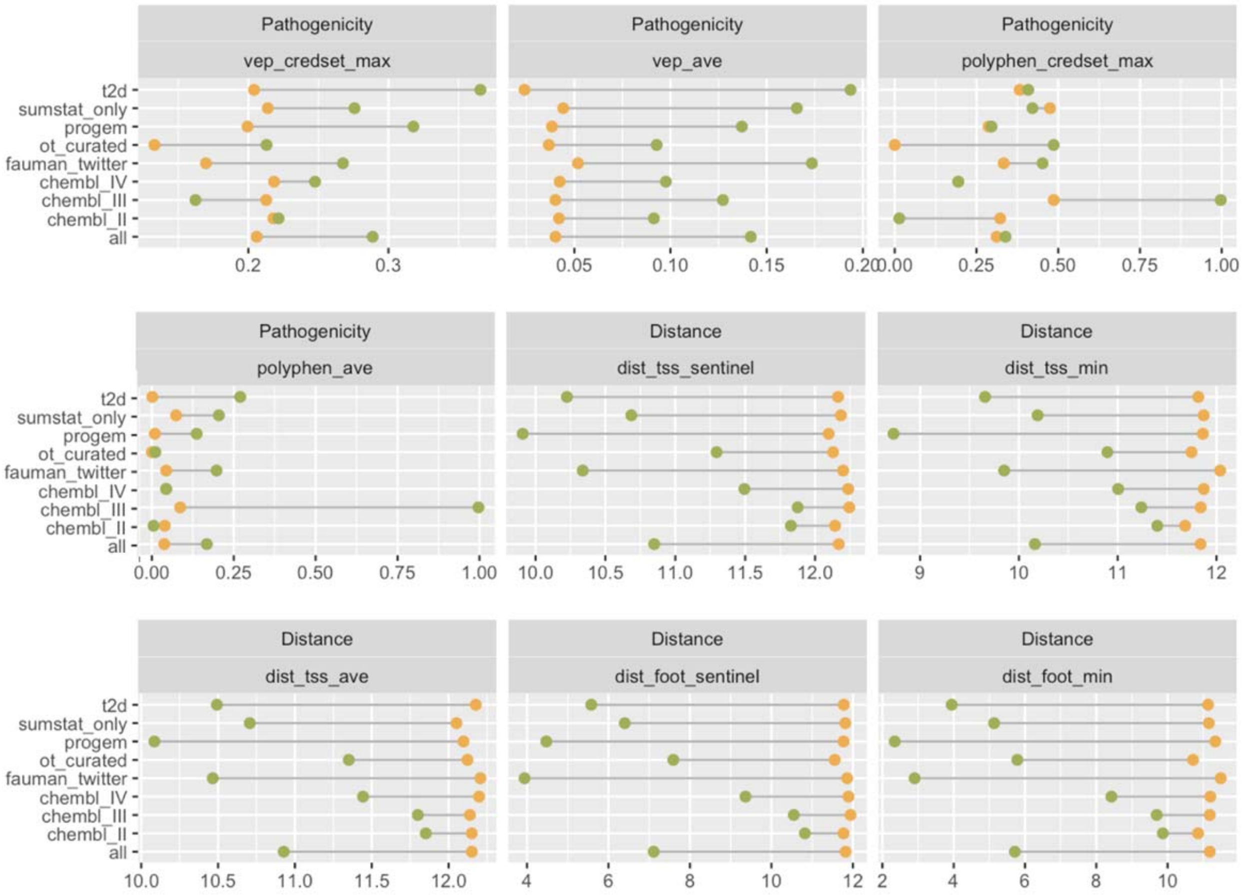


**Supplementary Figure 5**: Feature distributions. Each plot shows the mean value of a given predictor across different gold-standard datasets (y axis) for either gold standard positive genes (GSP, green) or gold standard negative genes (GSN, yellow). GSP genes are more easily distinguished from GSNs by distance in the manually curated datasets (especially Progem, Fauman_twitter, and T2D).
